# Gene expression signature for predicting homologous recombination deficiency in triple-negative breast cancer

**DOI:** 10.1101/2022.06.08.495296

**Authors:** Jia-Wern Pan, Zi-Ching Tan, Pei-Sze Ng, Muhammad Mamduh Ahmad Zabidi, Putri Nur Fatin, Jie-Ying Teo, Siti Norhidayu Hasan, Tania Islam, Li-Ying Teoh, Suniza Jamaris, Mee-Hoong See, Cheng-Har Yip, Pathmanathan Rajadurai, Lai-Meng Looi, Nur Aishah Mohd Taib, Oscar M. Rueda, Carlos Caldas, Suet-Feung Chin, Joanna Lim, Soo-Hwang Teo

## Abstract

Triple-negative breast cancers (TNBCs) are a subset of breast cancers that have remained difficult to treat. Roughly 1 in 10 of TNBCs arise in individuals with pathogenic variants in *BRCA1* or *BRCA2*, and treating BRCA-associated TNBCs with PARP inhibitors results in improved survival. A proportion of TNBCs arising in non-carriers of *BRCA* pathogenic variants have genomic features that are similar to *BRCA* carriers, and we postulated that gene expression may identify individuals with such features who might also benefit from PARP inhibitor treatment. Using genomic data from 129 TNBC samples from the Malaysian Breast Cancer (MyBrCa) cohort, we classified tumours as having high or low homologous recombination deficiency (HRD) and developed a gene expression-based machine learning classifier for HRD in TNBCs. The classifier identified samples with HRD mutational signature at an AUROC of 0.94 in the MyBrCa validation dataset, and strongly segregated HRD-associated genomic features in TNBCs from TCGA and METABRIC. Further validation of the classifier using the NanoString nCounter platform showed that the RNA-seq results correlated strongly with NanoString results (*r* = 0.90) from fresh frozen tissue as well as NanoString results from FFPE tissue (*r* = 0.84). Thus, our gene expression classifier may identify triple-negative breast cancer patients with homologous recombination deficiency, suggesting an alternative method to identify individuals who may benefit from treatment with PARP inhibitors or platinum chemotherapy.

**Novelty/Impact statement:** We developed a gene expression-based classifier for homologous recombination deficiency (HRD) in breast cancer patients using WES and RNA-seq data obtained from 129 TNBC samples from a Malaysian hospital-based cohort (MyBrCa). This classifier was able to predict for HRD status at an AUC of 0.94 in the MyBrCa cohort, and was also able to segregate HRD-associated features in TNBCs from TCGA. We also validated the classifier on a NanoString platform with both fresh frozen and FFPE tissue.

## Introduction

Triple negative breast cancer (TNBC) continues to be an area of unmet clinical need, as this aggressive subtype has fewer treatment options and higher mortality rates compared to other subtypes of breast cancer^1–3^. As a “catch-all” diagnosis for all breast tumours that test negative for both hormone receptors and HER2, TNBC is highly heterogeneous, and can be divided into several subtypes of its own^4,5^, each with potentially different treatment options. In this context, one potential biomarker that may be useful for subcategorizing TNBC patients for treatment is homologous recombination deficiency (HRD).

Cells with homologous recombination deficiency (HRD) have defects in their homologous recombination pathway leading to a diminished ability to repair DNA damage. In breast cancer, HRD is an important biomarker for therapies that utilize DNA-damaging agents such as platinum chemotherapy and PARP inhibitors. Platinum chemotherapies such as cisplatin, oxaliplatin, and carboplatin induce crosslinking of DNA that inhibits DNA repair and synthesis^6^. PARP inhibitors, on the other hand, are a new class of drugs that inhibit the action of Poly (ADP-ribose) polymerase (PARP) proteins, leading to double strand DNA breaks during cellular replication^7^. In tumours with HRD, these DNA-damaging agents cause an accumulation of mutations, leading to synthetic lethality and eventually cell death^8^. Tumours with deleterious genomic *BRCA* variants have defective homologous recombination repair pathways^9^, and PARP inhibitors have been approved for TNBC for individuals with deleterious germline variants in *BRCA1/2* (g*BRCA*m)^10,11^.

Besides deleterious germline *BRCA* variants, other molecular alterations in tumours may also lead to similar defects in the HRD pathway. This “BRCAness” feature has been proposed to broaden the patient population to PARP inhibitors^12^, and various molecular aspects of “BRCAness” have been characterized^13^. Some breast tumours with the “BRCAness” feature may arise when expressions of *BRCA1* or *BRCA2* are repressed by hypermethylation or somatic mutation^14^, or when the homologous recombination pathway is abrogated through mutations in other genes in the pathway (e.g. *PALB2* and *ATM*)^15,16^, and there are ongoing clinical studies that seek to expand the utility of PARP inhibitors in this context^17^. In addition, transcriptional signatures and genomic mutational signatures have been generated to identify tumours with HRD, with some association with PARP inhibitor sensitivity^18,19^. Clinical studies to examine the utility of these genomic signatures as predictive biomarkers have also been initiated^20^. In other cancer settings such as recurrent or high grade serous ovarian carcinoma, HRD assays such as the myChoice HRD assay and the Foundation Medicine T5 NGS assay have demonstrated some utility in guiding treatment with PARP inhibitors, and have received FDA approval as companion diagnostics^21–23^.

We have recently characterized the genomic and transcriptomic profiles of fresh frozen breast tumours from a large cohort of Malaysian patients of Chinese, Malay and Indian descent (the MyBrCa cohort)^24^. In order to study transcriptomic biomarkers for HRD in Asian TNBC, we first defined HRD status by clustering our TNBC samples using genomic features associated with HRD, followed by differential gene expression analyses comparing samples with high HRD to samples with low HRD. We identified a set of largely novel genes that were associated with HRD in our cohort, which we used to train a machine learning classifier to classify patient tumour samples as having high or low HRD. We validated the classifier using TNBC samples from the TCGA and METABRIC cohorts. We also validated the classifier on an alternative NanoString platform, using both FFPE and fresh frozen tumour samples. This classifier may have clinical utility as a non-gBRCAm biomarker to select for patients with high HRD who may benefit from treatment with platinum chemotherapy or PARP inhibitors.

## Methods

### Data description

Genomic sequencing data for this project was taken primarily from 94 TNBC samples included in the MyBrCa cohort tumour sequencing project. In brief, this included whole-exome sequencing (WES) and RNA-sequencing (RNA-seq) data collected from biobanked breast tumours of female patients from two hospitals – Subang Jaya Medical Centre in Subang Jaya, Malaysia, and Universiti Malaya Medical Centre in Kuala Lumpur, Malaysia, and analysed together with available clinical data. The cohort data and sequencing methods are described in full in Pan et al. (2020) and associated papers^24–26^. We also included an additional 35 TNBC samples that were not part of the original cohort description, for a total sample size of 129 MyBrCa TNBC samples. These samples were obtained and processed in largely the same way as the previous MyBrCa samples, with the only difference being the use of the Illumina NovoSeq 6000 as the sequencing platform instead of the Illumina HiSeq 4000. The sequencing coverage and quality statistics of WES and RNA-seq data for each new sample are summarized in Supplementary Table 1A and 1B, respectively. Additional validation data from TCGA and METABRIC TNBC samples were downloaded from the NIH Genomics Data Portal and the European Genome-phenome Archive, respectively.

### Transcriptomic data processing

Raw RNA-Seq reads were mapped to the hs37d5 reference human genome, and gene-level read counts were quantified using featureCounts (v. 1.2.31) with the Homo sapiens GRCh37.87 human transcriptome genome annotation model.

### Mutational analyses

To call SNVs, we used positions called by Mutect2 with following filters: minimum 10 reads in tumour and 5 reads in normal samples, OxoG metric less than 0.8, variant allele frequency (VAF) 0.075 or more, p-value for Fisher’s exact test on the strandedness of the reads 0.05 or more, and S_AF_ more than 0.75. For positions that are present in 5 samples or more, we removed two positions that were not in COSMIC and in single tandem repeats. We also removed variants that have VAF at least 0.01 in gnomAD, and considered only variants that are supported by at least 4 alternate reads, with at least 2 reads per strand. For indels, we also required the positions to be called by Strelka2. Variants were annotated using Oncotator version 1.9.9.0.

### Determination of HRD status

Genomic features from WES and sWGS data were used in a clustering step to group the TNBC samples into 2 groups: HRD high and HRD low. The genomic features used include telomeric allelic imbalance (TAI), loss of heterozygosity (LOH), large-scale transitions (LST), copy number amplification, copy number gain, copy number loss, copy number deletion, indel counts, and COSMIC mutational signature SBS3 scores. TAI, LOH and LST scores were determined using the scarHRD R package (v. 0.1.1)^27^ on allele-specific copy number profiles derived by Sequenza (v. 2.2) from paired tumour-matched normal WES bam files. The prevalence of the HRD-associated single base-pair substitution (SBS) mutational signature 3 from COSMIC (SBS3) was determined using deconstructSigs (v.1.8.0), restricted to samples with at least 15 SNVs. Scores for copy number amplification, gain, loss, and deletion were obtained using the QDNASeq R package (v. 1.22) on shallow-whole genome sequencing bam files. Scores for each feature were normalized using z-scores before clustering, except for indel counts which were log-transformed. K-means clustering and hierarchical clustering were performed using the Python packages “scikit-learn” (v. 1.2.1) and “scipy” (v. 1.5.2) respectively. Only samples that reached consensus between the two clustering algorithms were selected for further analysis, and the consensus clustering results were assigned as the HRD status of each sample.

### Differential expression analyses

Gene-level count matrices were normalised using the “Trimmed Mean of M-values” method implemented in the edgeR (v. 3.20.9) R package. The normalized count matrices were then transformed into log_2_ counts-per-million (CPM) values using the “cpm” function from the edgeR package in R. Differentially expressed genes were determined by empirical Bayes moderation of the standard errors towards a common value from a linear model fit of the transformed count matrices as implemented in the limma package, with the threshold for differential expression set as false discovery rate (FDR) < 0.001 and absolute log fold change >0.2.

### Pathway analysis

Over-representation analysis using KEGG and Reactome pathway-based sets as well as gene-ontology (GO) based sets was conducted using ConsensusPathDB (http://cpdb.molgen.mpg.de, accessed 21 April 2022) using the human database and ENSEMBL identifiers. For GO-based sets, the search was restricted to gene ontology level 2 and level 3 categories only.

Pathway analysis was conducted using gene set enrichment analysis (GSEA), as implemented in the Broad Institute GSEA Java executable (v 4.2.3), using the MSigDB Hallmark gene sets, as well as the KEGG gene sets, as implemented in the GSEA program using default options.

### Determination of germline *BRCA* mutation status

Carriers of deleterious pathogenic germline variants in *BRCA1* and *BRCA2* in the MyBrCa cohort were identified from targeted sequencing conducted as part of the BRIDGES study^28^. Each carrier was independently confirmed with Sanger sequencing.

### Classifier architecture

The machine learning framework was implemented in Python (v. 3.9.6) using the libraries “scikit-learn”, “scipy”, “numpy” (v. 1.19.2), “pandas” (v. 1.1.3). The input dataset consists of RNA-seq gene expression data quantified as TMM and log_2_ normalized counts per million (CPM), along with the HRD classification of each sample. The input data was split into training and validation sets following a 70/30 ratio using a 5-fold stratified shuffle split, resulting in five sets of data. The composite classifier combined two classifier pipelines for Support Vector Machine and Random Forest algorithms, respectively, and the final score (the probability that a sample is HRD High) was taken as the average score of both pipelines. The pipeline architecture was adapted from Sammut et al. (2021)^29^ and consists of four steps: collinearity removal, k-best feature selection, z-score scaling and classification. A randomized 1000-step 5-fold cross validation search that maximizes the area under the receiver operating characteristic (AUROC) curve is used to optimize all the hyperparameters for each classifier. This optimization is done separately for each of the five sets of training data, and each model is evaluated by taking the average of the AUROC scores produced by the 5-fold cross validation for the validation set. Of the five sets, the model with the highest average scores and the highest number of features used was selected as the final model for further validation (referred to below as the “MyBrCa model”).

### Validation on other cohorts

The classifier was validated using gene expression data from TNBC samples from other cohorts, including TCGA, the Molecular Taxonomy of Breast Cancer International Consortium (METABRIC) cohort^30,31^, and Nik-Zainal (2016)^32^ (NZ-560). Because the individual cohort datasets did not always contain all the genes used in the model training, the models used in each validation were retrained using the available genes for that cohort. The TCGA cohort RNA-seq data was downloaded from the GDC Data Portal and included all 217 genes used in the MyBrCa model. The METABRIC cohort, unlike our other cohorts, includes microarray data rather than RNA-seq data, and includes only 146 of the genes used in the MyBrCa model. Gene expression data for the METABRIC cohort was downloaded from the European Genome-phenome Archive. For the NZ-560 cohort, we used the log2 FPKM gene expression values from RNA-seq data that was reported in the original publication, but data was available for only 164 of the genes used in MyBrCa model. Gene expression values from each cohort were normalized using z-score scaling and quantile normalization separately for each cohort before classification.

### RNA extraction

RNA from tumour samples was extracted using the QIAGEN miRNeasy Mini Kit with a QIAcube, according to standard protocol. Total RNA was quantitated using a Nanodrop 2000 Spectrophotometer and RNA integrity was measured using an Agilent 2100 Bioanalyzer.

### NanoString validation

For the NanoString validation, we used data from a custom CodeSet developed for the NanoString nCounter platform. This custom CodeSet included 35 genes from our gene set and 3 housekeeping genes used for data normalization. We obtained NanoString nCounter read counts for these genes from 61 fresh frozen samples and 23 FFPE samples from the MyBrCa TNBC cohort. Expression for this gene set was measured on an nCounter MAX Analysis System, and the raw data was processed and normalized using the NanoString’s proprietary nSolver (v. 4.0) software before being exported as a normalized gene expression matrix text file for processing by the machine learning classifier, which was retrained using only the 35 genes included in the NanoString data. The NanoString gene expression values were normalized using z-score scaling and quantile normalization before classification.

### Statistical analyses

All box and whiskers plots in the figures are constructed with boxes indicating 25th percentile, median and 75th percentile, and whiskers showing the maximum and minimum values within 1.5 times the inter-quartile range from the edge of the box, with outliers not shown.

## Results

### Study population

Our discovery and training dataset consisted of 129 TNBC samples from the MyBrCa cohort, for which whole-exome sequencing and RNA-seq data were derived from fresh frozen primary tumour tissue and matched blood samples. 94 of these samples have been published previously (Pan et al. 2020), while the remaining 35 were sequenced more recently and have not been formally described. Of the 94 previously published samples, eight samples had pathogenic germline *BRCA1* variants, two samples had pathogenic germline *BRCA2* variants, and two samples had pathogenic germline *PALB2* variants^26^. The average age of the 129 TNBC patients was 52.9 years (± 13.6), and the majority of the samples were Stage II or Stage III, well-differentiated ductal carcinomas (Table 1). Our validation cohorts comprised of 87 TNBC tumours from The Cancer Genome Atlas (TCGA) breast cancer cohort^33^ (TCGA 2012), 306 TNBC tumour samples from the Molecular Taxonomy of Breast Cancer International Consortium (METABRIC) breast cancer cohort^30,31^, and 73 TNBC tumour samples from the Nik-Zainal (2016) breast cancer cohort (NZ-560)^32^.

**Table 1.** Clinico-pathological characteristics of the MyBrCaTNBC cohort. Statistical significance was calculated using student’s t-test for patient age and chi-square tests for all other variables, with N/A values excluded.

|  | MyBrCa TNBC Overall | MyBrCa TNBC HRD High | MyBrCa TNBC HRD Low | Statistical significance |
| --- | --- | --- | --- | --- |
| Subjects (n) | 129 | 41 | 72 |  |
| Patient age (yr ± SD ) | 52.9 ± 13.6 | 48.1 ± 11.7 | 56.5 ± 14.3 | p = 0.001 |
| **TNM stage (n(%))** |  |  |  | p = 0.83 |
| I | 20 (15.5) | 7 (17.1) | 11 (15.3) | p = 0.78 |
| II | 62 (48.1) | 22 (53.7) | 33 (45.8) | p = 0.39 |
| III | 41 (31.8) | 11 (26.8) | 25 (34.7) | p = 0.40 |
| IV | 2 (1.6) | 0 (0.0) | 2 (2.8) | p = 0.91 |
| N/A | 4 (3.1) | 1 (2.4) | 1 (1.4) |  |
| **Tumour Grade (n(%))** |  |  |  | p = 0.31 |
| 1 | 0 (0.0) | 0 (0.0) | 0 (0.0) |  |
| 2 | 20 (15.5) | 5 (12.2) | 13 (18.1) | p = 0.31 |
| 3 | 91 (70.5) | 33 (80.5) | 48 (66.7) | p = 0.31 |
| N/A | 18 (14.0) | 3 (7.3) | 11 (15.3) |  |
| **Histology subtype (n(%))** |  |  |  | p = 0.93 |
| Lobular Carcinoma | 2 (1.6) | 0 (0.0) | 2 (2.8) | p = 0.91 |
| Ductal Carcinoma | 106 (82.1) | 34 (82.9) | 60 (83.3) | p = 0.87 |
| Other | 7 (5.4) | 3 (7.3) | 4 (5.6) | p = 0.69 |
| N/A | 14 (10.9) | 4 (9.8) | 6 (8.3) |  |

### HRD in Asian TNBC

First, we examined the prevalence of HRD in the Asian TNBC setting by conducting an unsupervised clustering analysis of several genomic features associated with HRD in the MyBrCa TNBC samples. These features include commonly accepted features of HRD such as genomic loss of heterozygosity (LOH), telomeric allelic imbalance (TAI) and large-scale state transition (LST), short indels, copy number aberrations including gene amplification, gain, loss, and deletion, and the COSMIC mutational signature SBS3.

Hierarchical (Figure 1A) and k-means (Supp. Fig. 1) clustering of these genomic features revealed two distinct clusters which we called HRD High and HRD Low based on the levels of these HRD-associated scores. Out of the 129 samples that were clustered, 113 samples were concordant between the two clustering algorithms, while 16 samples were discordant and dropped from subsequent analyses. Of the 113 concordant samples, 41 (32%) were categorized as HRD High and 71 (68%) were categorized as HRD low.

**Figure 1.**
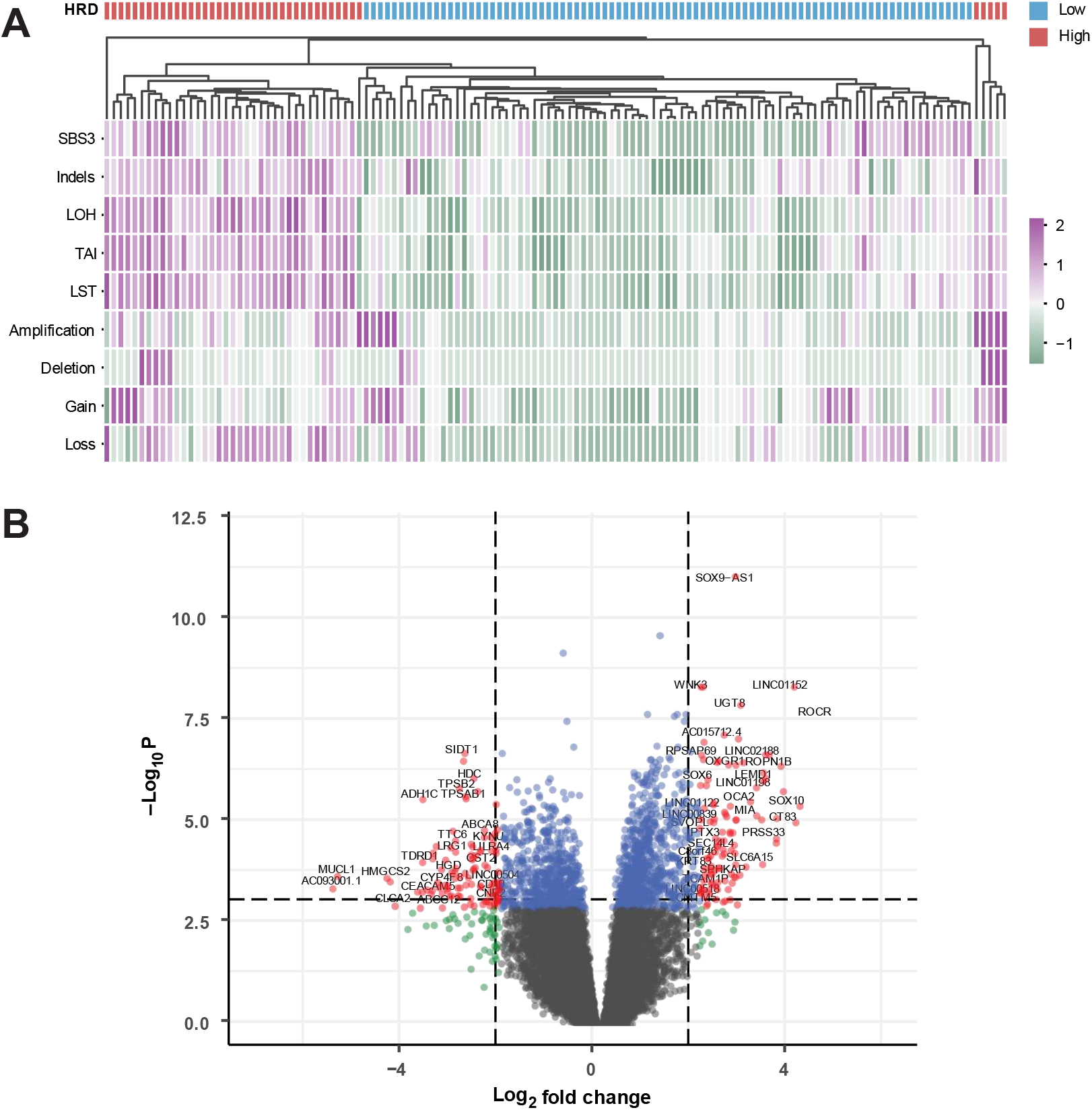
Clustering and gene expression analyses for homologous recombination deficiency (HRD) in MyBrCa TNBC samples. **(A)** Unsupervised hierarchical clustering of 129 MyBrCa TNBC samples using HRD-associated features including the COSMIC single base substitution mutational signature 3 (SBS3), short insertions and deletions (indels), loss-of-heterozygosity (LOH), telomeric allelic imbalance (TAI), large-scale transitions (LST), as well as copy number amplifications, deletions, gain and loss. All scores were scaled using z-scores, and the indels score was also log-transformed prior to scaling esignati ns f eac sa le as ig w a e indicated by t e “ “ ann tati n bar. **(B)** Volcano plot for differential expression analysis comparing HRD High and HRD Low samples. Dotted lines indicate the thresholds used to classify genes as differentially expressed (Benjamini-Hochberg adjusted p-value < 0.001, absolute log_2_ fold change > 2).

Next, using differential gene expression analysis of RNA-sequencing data from the TNBC tumour samples, we identified a set of 217 genes that were differentially expressed between the two groups by manually curating the top upregulated and downregulated genes based on Benjamini-Hochberg adjusted p-values of less than 0.001, with a minimum log_2_ fold change of 2 in either direction (Figure 1B, Supp. Table 2). This gene set of 217 genes was substantially different from previous HRD-associated gene sets derived in studies by Peng et al. 2014^34^and the i-SPY 2 consortium^35,36^, with very small numbers of overlapping genes (1 between MyBrCa and Peng, 15 between MyBrCa and i-SPY 2; Supp. Fig. 2).

Over-representation analysis of the selected 217 differentially expressed genes using KEGG and Reactome pathway-based gene sets revealed an enrichment in a number of metabolic and signaling pathways (Supp. Table 3), but not pathways known to be associated with HRD. Similarly, analysis of over-represented gene ontology (GO) terms for the 217 genes showed an enrichment of extracellular matrix, plasma membrane, and hormone-related terms, but not HRD-related terms (Supp. Table 4), suggesting that the 217 genes were largely not previously known to be associated with HRD. On the other hand, gene set enrichment analysis (GSEA) of MSigDB Hallmark and KEGG pathways comparing TNBC tumour samples classified as HRD High to samples classified as HRD Low revealed an upregulation of cell cycle and DNA repair pathways in samples classified as HRD High, suggesting that HRD pathways are indeed differentially expressed between the two groups when looking across the whole transcriptome (Supp. Table 5-6). Interestingly, immune-related pathways appear to be downregulated in the HRD High group and upregulated in the HRD Low group as well (Supp. Table 5-6).

### Classification of tumour samples according to HRD status using the gene set

Using the set of 217 genes identified above, we trained a machine learning classifier, which we call HRD200, to classify the HRD status of any given tumour sample, using an adaptation of the composite classifier framework described in Sammut et al. (2021)^29^. Our adaptation of this framework incorporates 5-fold stratified shuffling to split the samples along 70/30 ratio for model training and testing, respectively, followed by feature selection and 5-fold cross-validation (see Methods). The best model under this framework achieved a mean AUROC of 0.91 in training and 0.94 on the test dataset, utilizing 169 of the 217 genes as features (Figure 2A). Overall, the HRD200 classifier incorporating the best model designated 34 out of the 41 Asian TNBC in the training samples with high HRD scores correctly as “HRD High” and classified 62 out of 72 of the samples with low HRD scores correctly as “HRD Low”, for an F1 score of 0.80 and an accuracy of 85%. The classifier also categorized 5 out of the 7 samples in our cohort with known pathogenic germline BRCA1 variants as HRD High (Figure 2B). In addition, the use of our specific 217 gene set with this classification approach was able to outperform similar classifiers using the gene sets by Peng and i-SPY 2, although the difference was small (Supp. Figure 3).

**Figure 2.**
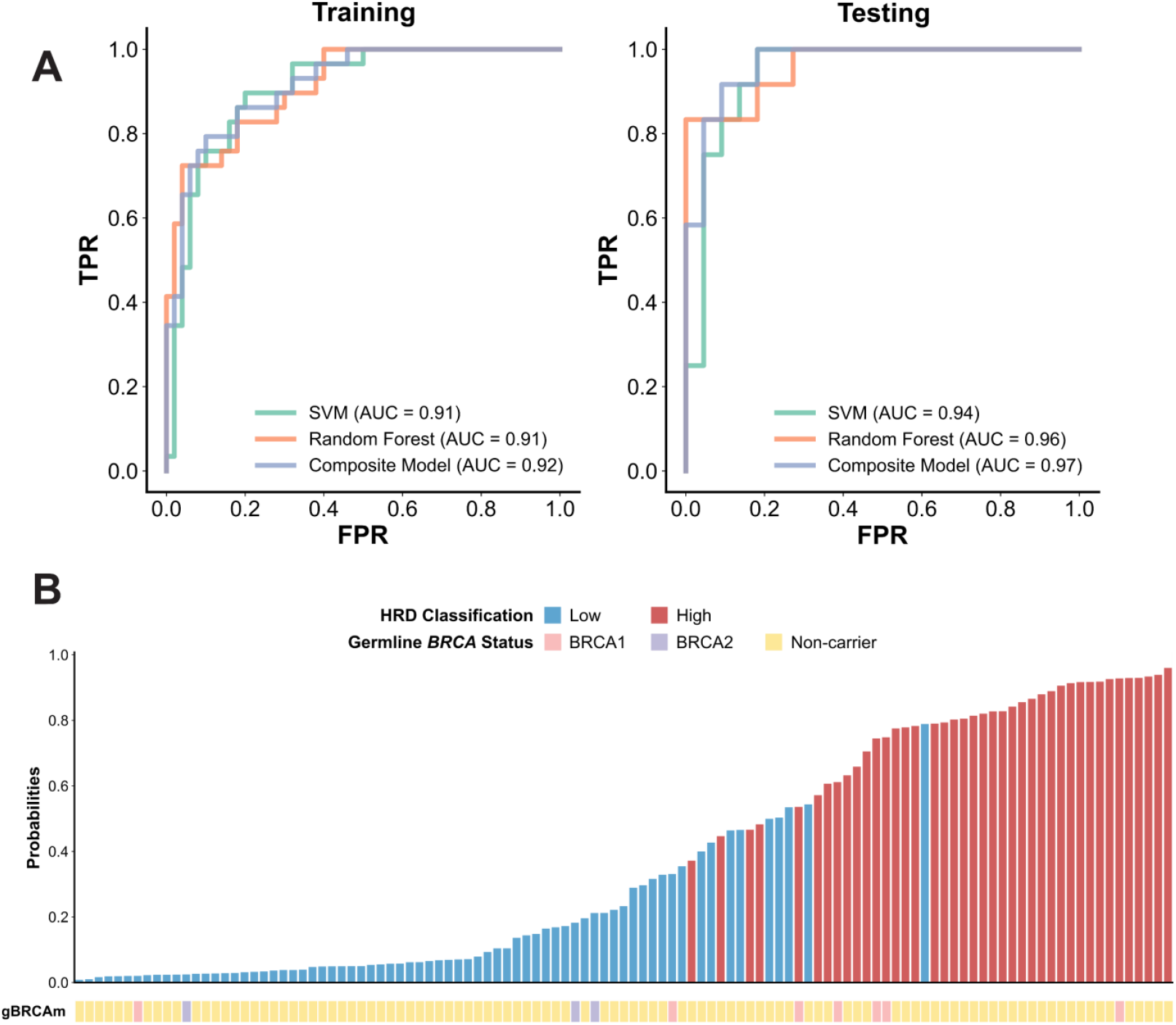
Performance of the HRD200 classifier in the MyBrCa TNBC cohort. **(A)** Receiver operating characteristic (ROC) curves of false positive rate (FPR) and true positive rate (TPR) showing the performance of the HRD200 composite classifier in predicting HRD High status in 70:30 training:testing gene expression datasets from 113 MyBrCa TNBC samples. The HRD200 classifier was trained on gene expression data of 217 differentially expressed genes. The ROC curves for the support vector machine (SVM), random forest (RF), and composite models are shown separately. (B) Bar chart showing the probability of a sample being HRD High according to the HRD200 classifier, compared to their ground-truth HRD classification (color of the bar) and known germline *BRCA* status (gBRCAm annotation).

### Validation of the HRD200 classifier in other cohorts

Next, we examined the ability of our HRD200 classifier to predict HRD-associated genomic features in other cohorts. First, we used our classifier to predict the HRD status of 87 TNBC tumours in The Cancer Genome Atlas (TCGA) dataset (TCGA 2012) for which mutational signature data was available. Of the 87 samples, our classifier designated 56 of these samples as HRD High and 31 as HRD Low. A comparison of these two groups revealed that the HRD High samples had significantly higher levels of the HRD-associated SBS3 mutational signature, as well as higher numbers of indels and copy number aberrations (Figure 3), suggesting that our classifier was able to successfully segregate samples with HRD-associated features in the TCGA TNBC cohort. We also tested our classifier on 306 TNBC samples from the METABRIC cohort and 73 samples from the Nik-Zainal (2016) cohort, although in both cases we had to reduce the number of genes included in the classifier as expression data was not available for some genes. In both the METABRIC and Nik-Zainal cohorts, samples designated as HRD High by our classifier had significantly higher levels of the SBS3 mutational signature, and the HRD High METABRIC samples also had significantly more copy number aberrations (Supp. Figure 4).

**Figure 3.**
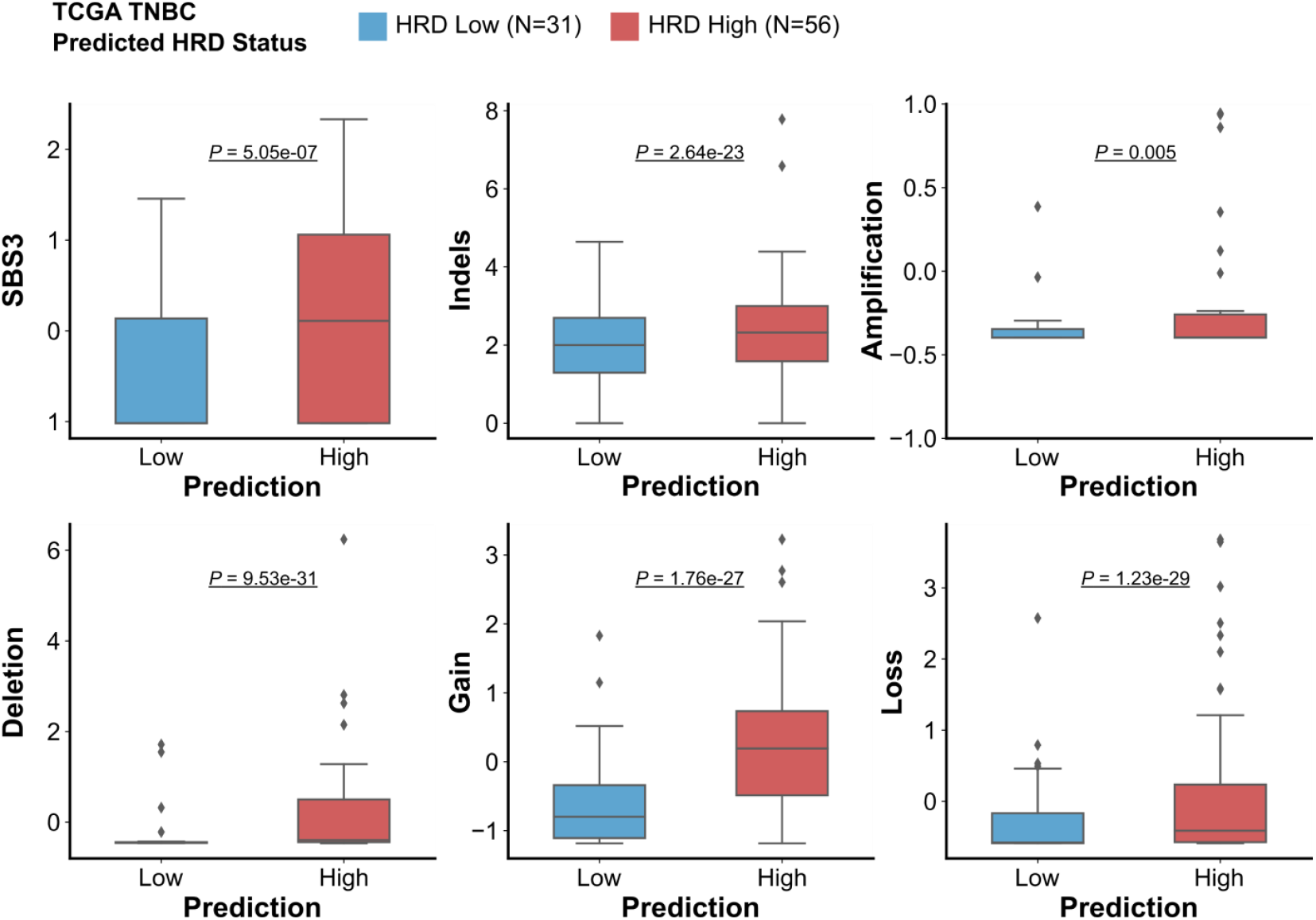
Validation of the HRD200 classifier in the TCGA cohort. Comparisons of COSMIC single base substitution mutational signature 3 (SBS3), short insertions and deletions (indels), copy number amplifications, deletions, gain and loss between TCGA TNBC samples classified by the HRD200 classifier as HRD Low (n=31, in blue) and samples classified as HRD High (n=56, in red). All scores were scaled using z-scores, and the indels score was also log-transformed prior to scaling.

### Validation of the HRD classifier using a NanoString platform

Next, we evaluated the robustness of the HRD200 classifier with regards to different methods of measuring gene expression, as well as to the use of formalin-fixed paraffin-embedded (FFPE) tissue rather than fresh frozen tissue. We did this because both RNA-seq data as well as fresh frozen tumour tissue are expensive and difficult to obtain as part of routine clinical practice.

To evaluate the performance of the HRD200 classifier with a different gene expression measurement method, we used data from the NanoString nCounter platform^37^, which uses direct digital detection of mRNA molecules to generate gene-level transcript counts. We were able to obtain gene expression data for 35 genes of our gene set from 61 fresh frozen tissue samples as well as 23 FFPE tissue samples from the same cohort of MyBrCa TNBC patients described above, using a custom NanoString nCounter CodeSet. These data were then inputted into a version of our HRD200 classifier that was optimized for the 35 genes, and the results were compared to the HRD200 classification results from the original RNA-seq data. Using the original RNA-seq classifications as the ground truth, the NanoString-based classification had an AUROC of 0.97 for fresh frozen tissue and 0.98 for FFPE (Figure 4A). Additionally, the probabilities for any given sample being classified as HRD High from RNAseq and NanoString data were highly correlated, with a Spearman’s correlation coefficient (ρ) of 0.9 when comparing RNASeq and fresh frozen NanoString results, and a (ρ) of 0.84 when comparing RNAseq and FFPE NanoString results (Figure 4B). Pairwise comparisons of the RNAseq and NanoString results, again treating RNAseq data as the ground truth, revealed an overall concordance of 0.93 and an F1 score of 0.90 for fresh frozen tissue, and a concordance of 0.89 and an F1 score of 0.86 for FFPE tissue (Figure 4C, Supp. Fig. 5). Overall, this suggests that our HRD200 classifier was robust even when used with expression data from a different platform, and also when used with smaller subsets of genes compared to the original gene set.

**Figure 4.**
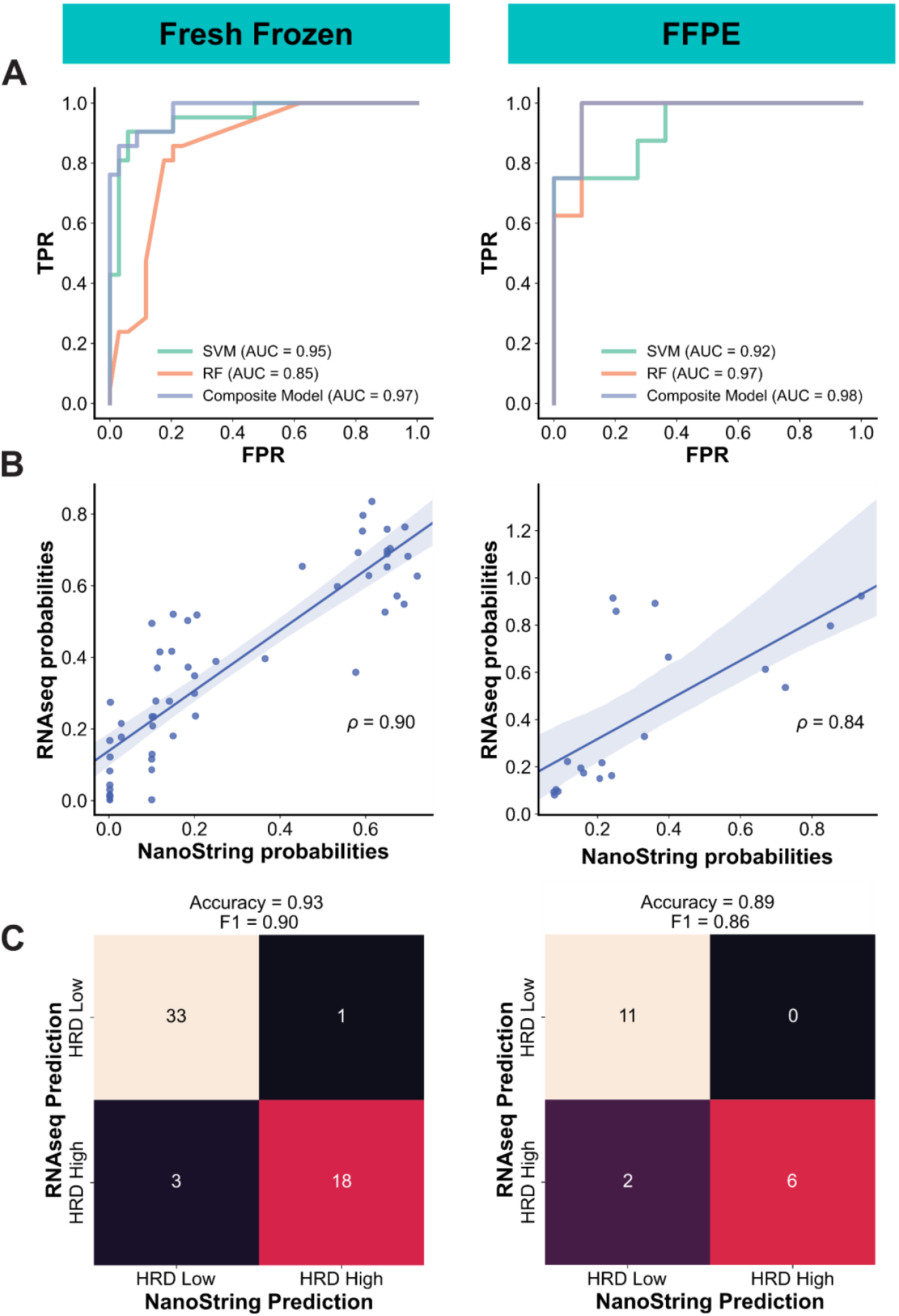
Validation of the HRD classifier on the NanoString nCounter platform. **(A)** Receiver operating characteristic (ROC) curves showing the performance of HRD200 classifier (retrained using a 35 gene subset) in predicting HRD High status from NanoString nCounter gene expression data from fresh frozen (left, n=61) and FFPE (right, n=23) samples from the MyBrCa TNBC cohort, using HRD High predictions from RNAseq data by the HRD200 classifier as the ground truth. ROC curves shown are for the testing dataset from 70:30 training:testing splits of the samples. The ROC curves for the support vector machine (SVM), random forest (RF), and composite models are shown separately. **(B)** Comparison of the HRD High probabilities given by the HRD200 classifier for RNAseq (y-axis) and NanoString (x-axis) fresh frozen (left) or FFPE (right) matched samples. Also shown are Spearman’s correlation coefficient (ρ) for each comparison. **(C)** Confusion matrices comparing HRD200 classification of samples using RNAseq data to classification using NanoString data from fresh frozen (left) or FFPE (right) samples.

## Discussion

Here, we describe the development and validation of a gene expression classifier for homologous recombination deficiency in Asian TNBC. Using mutation load and other genomic features, we were able to sort our TNBC samples into two clusters - HRD high and HRD low, and we derived a gene set associated with HRD status using gene expression analyses. Using that gene set, we subsequently developed an ensemble machine learning model classifier (HRD200) that could discriminate between samples with high versus low HRD with good accuracy and AUROC from gene expression data. The classifier also had the ability to segregate samples according to genomic features associated with HRD in TNBC validation cohorts from TCGA, METABRIC, and Nik-Zainal (2016). Importantly, we also found a very high concordance rate in the classification results when using an alternative measure of gene expression (NanoString), as well as when using FFPE material instead of fresh frozen tissue, even with small subsets of the gene panel, suggesting that the HRD200 classifier could be robust for real-world clinical use situations.

In the TNBC setting, we found a large cluster of samples with high HRD-associated genomic features and mutational signatures. These results suggest that HRD is a significant driver of tumour mutations in Asian TNBC, even in the absence of pathogenic germline *BRCA* variants. This in turn suggests that there may be a significant number of Asian TNBC patients lacking pathogenic germline *BRCA* variants who may still benefit from therapies that target the HRD pathway, such as platinum chemotherapy or PARP inhibitors. Thus, tools to detect HRD in tumour samples may have clinical utility as a non-gBRCAm biomarker to select for Asian breast cancer patients with high HRD who may benefit from treatment with platinum chemotherapy or PARP inhibitors.

The training and validation of the HRD classification tool described in this paper relies in part on measures of the mutational signature SBS3 in each tumour sample. The COSMIC mutational signature SBS3 is a well-studied cancer mutational signature that has been validated in orthogonal techniques^38,39^, and its HRD-driven etiology has been experimentally confirmed^40^. The use of HRD-associated mutational signatures to predict “BRCAness” and thus response to PARP inhibitors and platinum chemotherapy has received significant attention in recent years, with tools such as HRDetect^19^ and CHORD^41^ demonstrating that HRD-associated mutational signatures can be used to predict BRCA1/BRCA2-deficiency in breast tumours as well as survival of breast cancer patients^42^. The clinical utility of such tools in a breast cancer setting is being tested in ongoing clinical trials; however, one significant drawback is that these methods usually require whole-genome sequencing of tumour samples, which may be prohibitively expensive and time-consuming, particularly in low-resource settings. By using gene expression signatures, our method for predicting HRD in tumour samples may offer a useful alternative.

As mentioned above, other gene expression signatures related to HRD have also been previously described - Peng and colleagues (2014)^34^ described a set of 230 genes associated with homologous recombination DNA repair in a microarray analysis of a nonmalignant human mammary epithelial cell line that predicts clinical outcome in cancer patients, while a different 77-gene panel of “*BRCA1*ness” was significantly associated with response to PARP inhibitor treatment in the I-SPY 2 breast cancer clinical trial^35,36^. Interestingly, there appears to be little overlap between the different gene panels – only one out of the 217 genes in our gene set are included in Peng’s 230-gene set, and only 15 of the 217 genes in our study are included in the 77-gene “BRCA1ness” panel, with zero genes common across all three sets. The lack of overlap may reflect differences in methodology or study population, and further study to reconcile these differences may be warranted. Nonetheless, when the same machine learning approach was used, all three gene sets were almost equally predictive for HRD status in our dataset, with very high predictive value across the board. This suggests that the machine learning methodology used may be more important than the specific gene set for researchers to be able to derive accurate predictions of HRD status from gene expression data, as long as the gene set used retains some association with HRD in the study population.

Taken together, we believe that the HRD200 classifier, implemented as a NanoString-based test, may have clinical utility as a non-*BRCA*m biomarker to select for patients with high HRD who may benefit from treatment with PARP inhibitors. However, further development of the classifier will require the initiation of clinical trials in order to determine if HRD200 can correctly identify patients who are sensitive to PARP inhibitor therapy in real-world clinical settings.

## Abbreviations

FFPE: formalin-fixed, paraffin-embedded
g*BRCA*m: germline *BRCA1*/*BRCA2* mutations
HRD: homologous recombination deficiency
MyBrCa: the Malaysian Breast Cancer cohort
PARP: Poly (ADP-ribose) polymerase
RNA-seq: RNA sequencing
SBS3: COSMIC single base-pair substitution mutational signature 3
TCGA: The Cancer Genome Atlas
TNBC: triple-negative breast cancer
WES: whole-exome sequencing

## Acknowledgements

Cancer Research Malaysia receives charitable funding from the Scientex Foundation, Estée Lauder Companies, Vistage Malaysia, Yayasan PETRONAS, and Yayasan Sime Darby which contributed to the funding of this study. Funding was also provided by a research grant from the Newton-Ungku Omar Fund (MRC Ref: MR/P012442/1) to SFC and SHT. OMR, CC, and SFC also receive funding from Cancer Research UK. All genomics work was undertaken by the Genomics Core Facility CRUK Cambridge Institute.

## Author Contributions

JWP led the data analysis, supervised experiments, and wrote the manuscript. ZCT, MMAZ and PNF contributed to data analysis and generation of figures. PSN contributed to data analysis, experimental design, sample collection, and carried out experiments. JYT, SNH, TI, LYT, SJ, MHS, CHY, PR, LML, and AMT contributed to sample collection and processing and data collection, while OMR and SFC generated and processed sequencing data. PR and LML provided histopathology expertise, and collected clinical data together with MHS, TI, SJ, LYT, CHY and AMT. OMR, CC, and SFC also interpreted results and helped to draft the manuscript. CC and SHT contributed to obtaining funding for the project. SHT also designed experiments, drafted the manuscript, and provided overall project direction and supervision. The work reported in the paper has been performed by the authors, unless clearly specified in the text.

## Data Accessibility

The WES and RNA-seq data generated in this study are available in the European Genome-phenome Archive under accession number EGAS00001006518. Previously published data from Pan et al. (2020) are available in EGA under accession numbers EGAS00001004518. Access to controlled patient data will require the approval of the Data Access Committee. Further information is available from the corresponding author upon request.

## Ethics Declaration

Patient recruitment and sample collection was reviewed and approved by the Independent Ethics Committee, Ramsay Sime Darby Health Care (reference no: 201109.4 and 201208.1), as well as the Medical Ethics Committee of the University Malaya Medical Centre (reference no: 842.9). Written informed consent to participation in research was given by each individual patient.

---

**Supp. Fig. 1.**
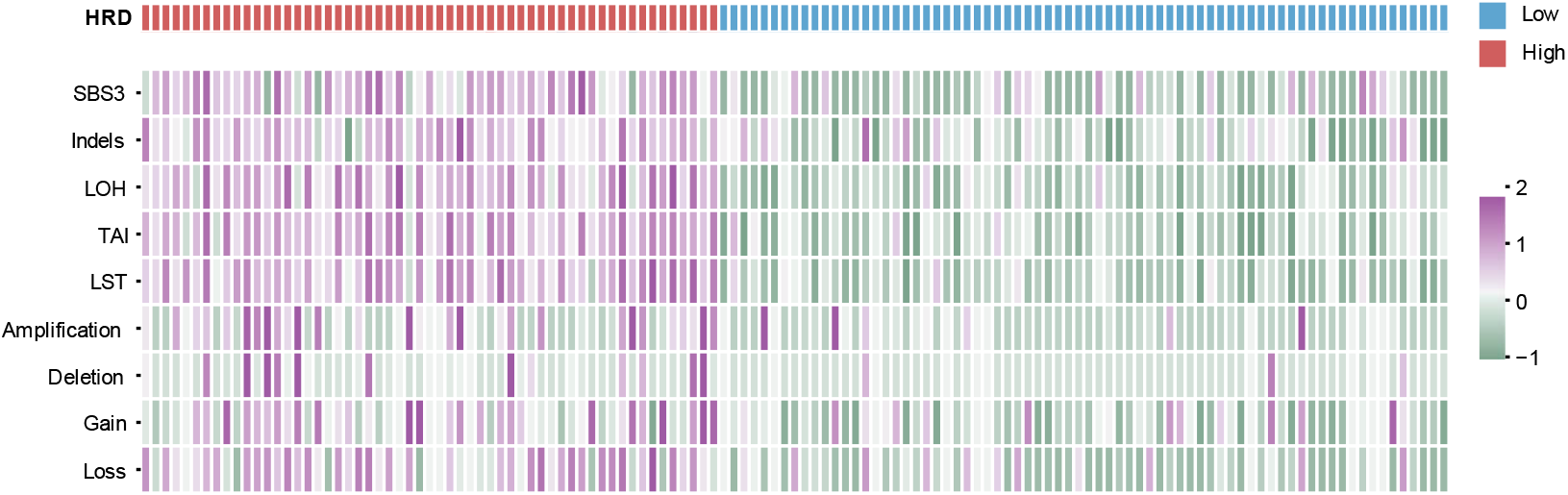
K-means clustering of 129 MyBrCa TNBC samples using HRD-associated features. Features used include the COSMIC single base substitution mutational signature 3 (SBS3), short insertions and deletions (indels), loss-of-heterozygosity (LOH), telomeric allelic imbalance (TAI), large-scale transitions (LST), as well as copy number amplifications, deletions, gain and loss. Clustering was performed using *k* = 2. All scores were scaled using z-scores, and the indels score was also log-transformed prior to scaling. Designations for each sample as HRD High or HRD Low are indicated by the “HRD” annotation bar.

**Supp. Fig. 2.**
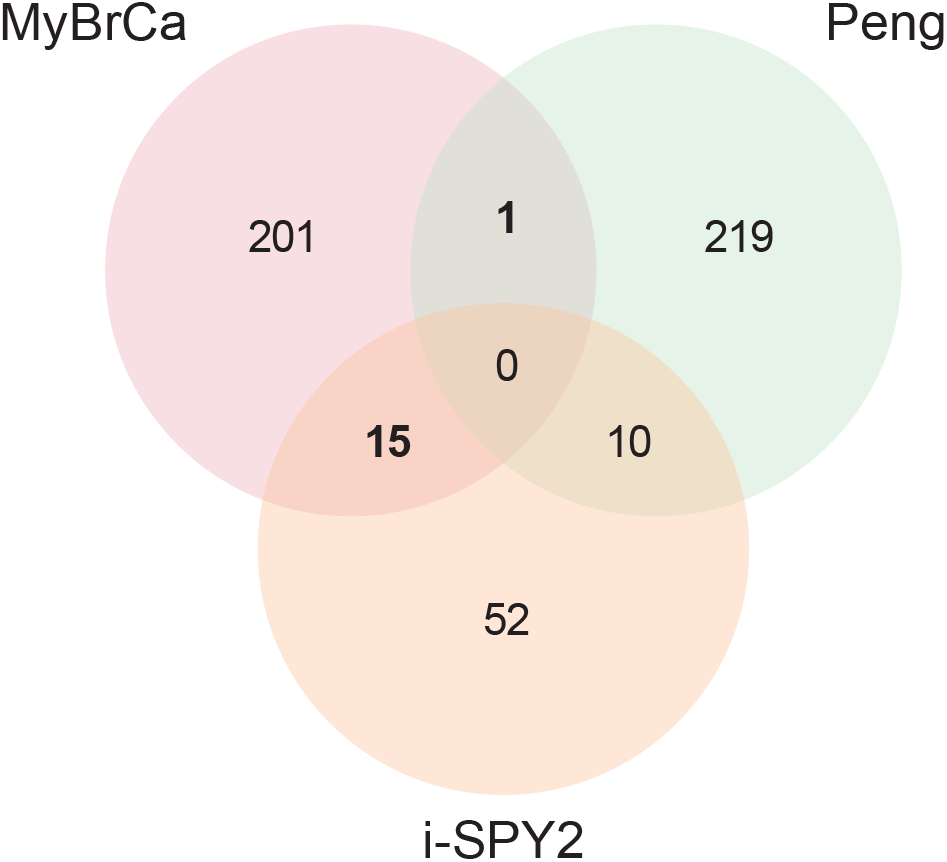
Venn diagram of overlapping genes in the MyBrCa (this study), Peng (2014) and i-SPY2 gene sets indicative of HRD.

**Supp. Fig. 3.**
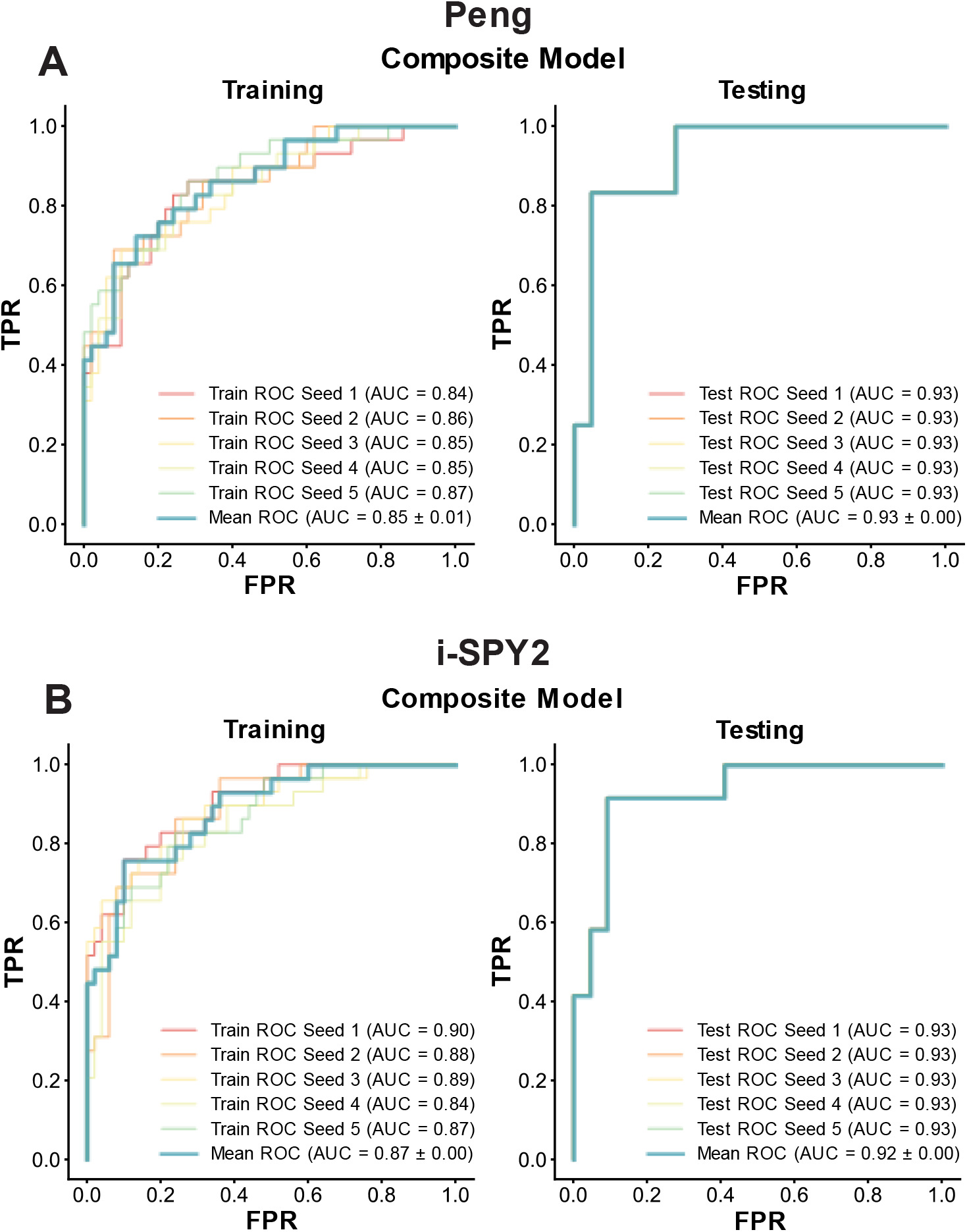
Performance of composite models using the Peng (2014) and i-SPY2 gene sets in predicting HRD High status in the MyBrCa TNBC cohort. Shown are the receiver operating characteristic (ROC) curves of false positive rate (FPR) and true positive rate (TPR) showing the performance of the composite classifiers trained using the Peng (2014) gene set of 230 HRD-associated genes (A) as well as the i-SPY2 gene set of 77 HRD-associated genes (B) across 5 cross-validation seeds in 70:30 training:testing gene expression datasets from 113 MyBrCa TNBC samples. The mean ROC across the 5 seeds are also shown.

**Supp. Figure 4.**
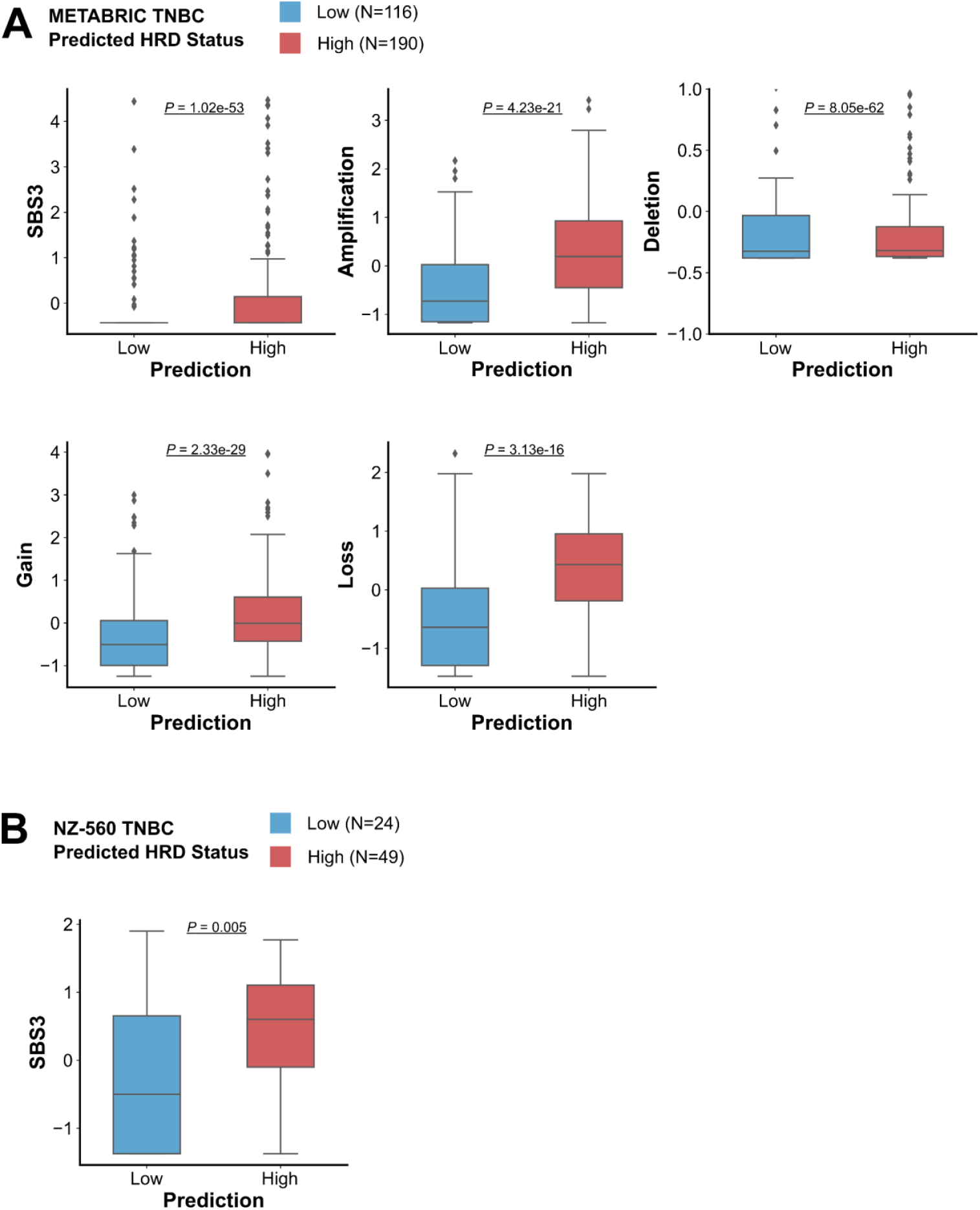
Validation of the HRD classifier in the METABRIC and NZ-560 cohorts. **(A)** Comparisons of COSMIC single base substitution mutational signature 3 (SBS3), copy number amplifications, deletions, gain and loss between METABRIC TNBC samples classified by the HRD200 classifier (retrained using 146 genes) as HRD Low (n=116, in blue) and samples classified as HRD High (n=190, in red). **(B)** Comparison of SBS3 between NZ-560 TNBC samples classified by the HRD200 classifier (retrained using 164 genes) as HRD Low (n=24, in blue) and samples classified as HRD High (n=49, in red). All scores were scaled using z-scores, and the indels score was also log-transformed prior to scaling.

**Supp. Fig. 5.**
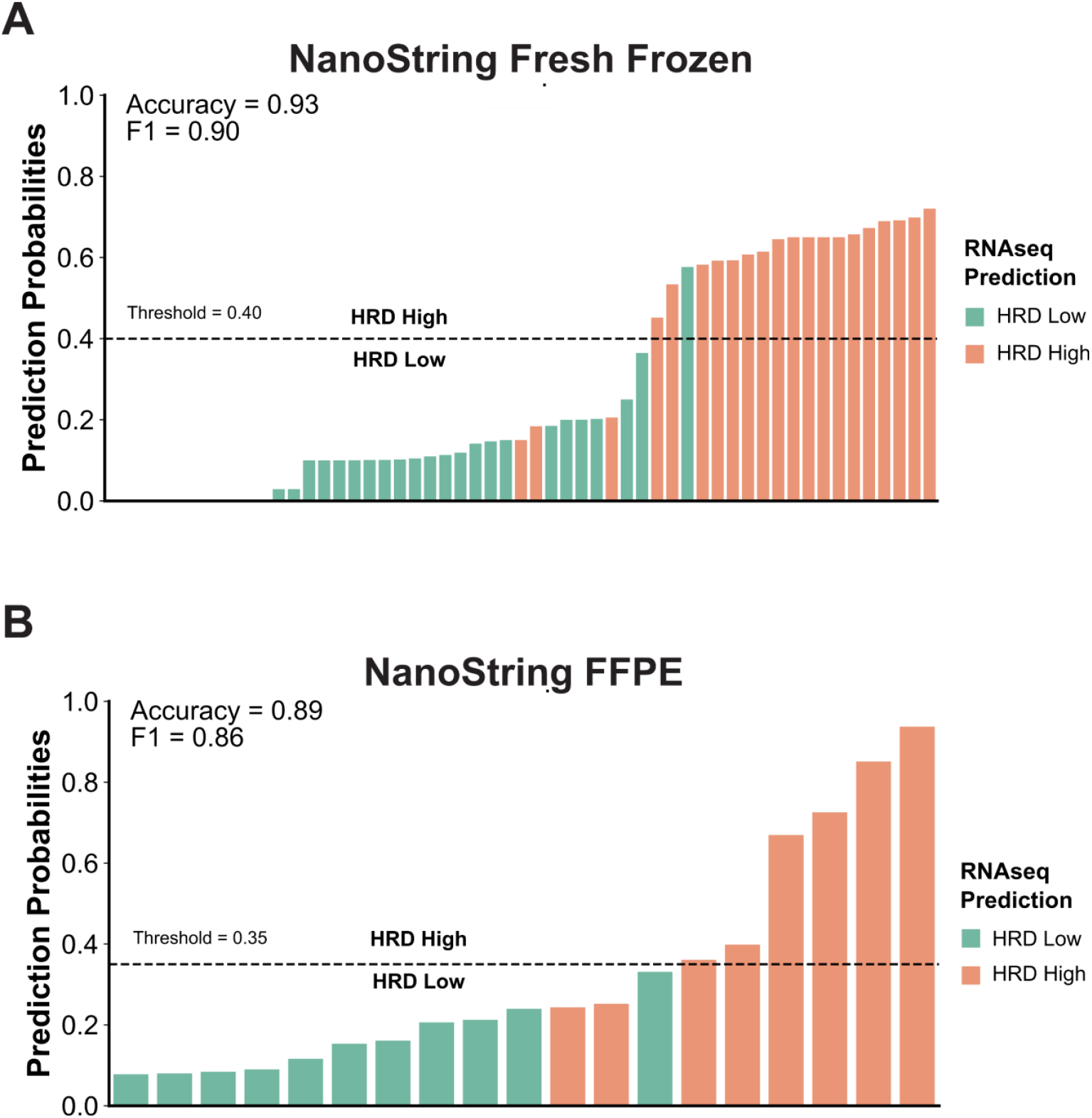
Comparison of RNAseq and NanoString predictions. Bar charts showing the probability of a sample being HRD High according to the HRD200 classifier using NanoString nCounter data from (A) fresh frozen (n=61) and (B) FFPE samples (n=23) from the MyBrCa TNBC cohort, compared to their HRD classification from RNAseq data (color of the bar). The probability threshold used for HRD classification of the NanoString data is indicated with a dotted line.

